# A closer look at high-energy X-ray-induced bubble formation during soft tissue imaging

**DOI:** 10.1101/2023.02.14.528474

**Authors:** R. Patrick Xian, Joseph Brunet, Yuze Huang, Willi L. Wagner, Peter D. Lee, Paul Tafforeau, Claire L. Walsh

**Author notes:** **Corresponding authors:** Paul Tafforeau, Peter D. Lee, R. Patrick Xian. These authors contributed equally to the work (R. Patrick Xian, Joseph Brunet). **Author Contributions:** R.P.X., J.B., P.D.L., P.T., C.L.W. conceptualized the project and designed experiments. R.P.X., J.B., P.T., C.L.W performed the sample preparation and imaging. R.P.X., J.B., Y.H., W.L.W., P.D.L., P.T., C.L.W. analyzed the data. R.P.X., J.B., P.D.L., P.T., C.L.W. wrote the paper.

## Abstract

Improving the scalability of tissue imaging throughput with bright, coherent X-rays requires identifying and mitigating artifacts resulting from the interactions between X-rays and matter. At synchrotron sources, long-term imaging of soft tissues in solution can result in gas bubble formation or cavitation, which dramatically compromises image quality and integrity of the samples. By combining in-line phase-contrast cineradiography with *operando* gas chromatography, we were able to track the onset and evolution of high-energy X-ray-induced gas bubbles in ethanol-embedded soft tissue samples for tens of minutes (2 to 3 times the typical scan times). We demonstrate quantitatively that vacuum degassing of the sample during preparation can significantly delay bubble formation, offering up to a twofold improvement in dose tolerance, depending on the tissue type. However, once nucleated, bubble growth is faster in degassed than undegassed samples, indicating their distinct metastable states at bubble onset. Gas chromatography analysis shows increased solvent vaporization concurrent with bubble formation, yet the quantities of dissolved gases remain unchanged. Coupling features extracted from the radiographs with computational analysis of bubble characteristics, we uncover dose-controlled kinetics and nucleation site-specific growth. These hallmark signatures provide quantitative constraints on the driving mechanisms of bubble formation and growth. Overall, the observations highlight bubble formation as a critical, yet often overlooked hurdle in upscaling X-ray imaging for biological tissues and soft materials and we offer an empirical foundation for their understanding and imaging protocol optimization. More importantly, our approaches establish a top-down scheme to decipher the complex, multiscale radiation-matter interactions in these applications.

**Significance statement:** Better probing the X-ray radiation dose limit of bubble formation in biological tissue and developing mitigation methods is essential for improving imaging techniques involving X-ray, such as synchrotron X-ray tomography or crystallography. Here, we combined *operando* gas chromatography with in-line X-ray phase-contrast radiography on human lung and brain tissue to investigate bubble formation under high-energy X-ray irradiation. We demonstrate that vacuum degassing delays bubble nucleation up to a factor two, depending on the tissue type. Gas chromatography analysis showed increased solvent vaporization during bubble formation; however, the quantities of dissolved gases remained unchanged. Moreover, depending on the nucleation site, bubble growth can be geometrically constrained by sample microstructure, which influence its dynamics.

## Introduction

X-ray imaging and spectroscopic techniques are being gradually adopted and refined for probing biological systems with unprecedented resolution and sensitivity[1], [2]. Soft biological samples are prone to radiation damage[3], [4] in their natural conditions or the preferred condition to study, which requires procedural adjustment of existing X-ray techniques which are more attuned to studying crystalline samples or small samples of (radiation) hard materials. For X-ray imaging, wet embedding in liquid solvents or gels is inherently an efficient and scalable preparation method because it can accommodate deformable samples of arbitrary sizes, from thin or thick slices[5] to entire organs[6] and whole organisms[7]–[9]. This compares favorably with the use of solid paraffin or resin for sample immobilization[10], which are slow to impregnate thick tissues, or high-pressure freezing[11], which is currently limited to submillimeter-thin samples. In imaging wet biological samples embedded in solution or gel, the formation of gas bubbles and their dynamical evolution are important issues that compromise the outcome[12], [13], but to date lack quantitative study. Bubbles in this context are closed gas-liquid interfaces exhibiting a high refractive index gradient across the boundary[14]. For X-ray phase contrast, often adopted for imaging soft tissues[6], [15]– [17], the strong edge-enhancing effect from gas bubbles[18], [19] often largely eclipses the inherent phase contrast from (unstained) soft tissues[20]. Although exogenous micrometer-sized bubbles (microbubbles) may be injected into the sample to enhance contrast[21], [22], their uncontrolled creation during X-ray irradiation[23] is detrimental to long scans often required for high-resolution imaging of large samples[6] or for *in vivo* dynamic monitoring in developmental biology[8] and physiology[24]. Due to bubble growth and their motion, the experiments need to be interrupted and the sample reprocessed to mitigate the strong imaging artifacts[25] (see Fig. 1a-c). The cavitation process and the subsequent bubble motion can potentially also cause damage[26], [27] to fragile tissue microstructures, like brains or embryos[8], especially in high-resolution bioimaging settings that are increasingly being adopted for elucidating multiscale information and spatiotemporal processes. Therefore, identification of the experimental conditions and underlying mechanisms that induce bubble formation in high energy X-ray imaging is crucial.

**Figure 1.**
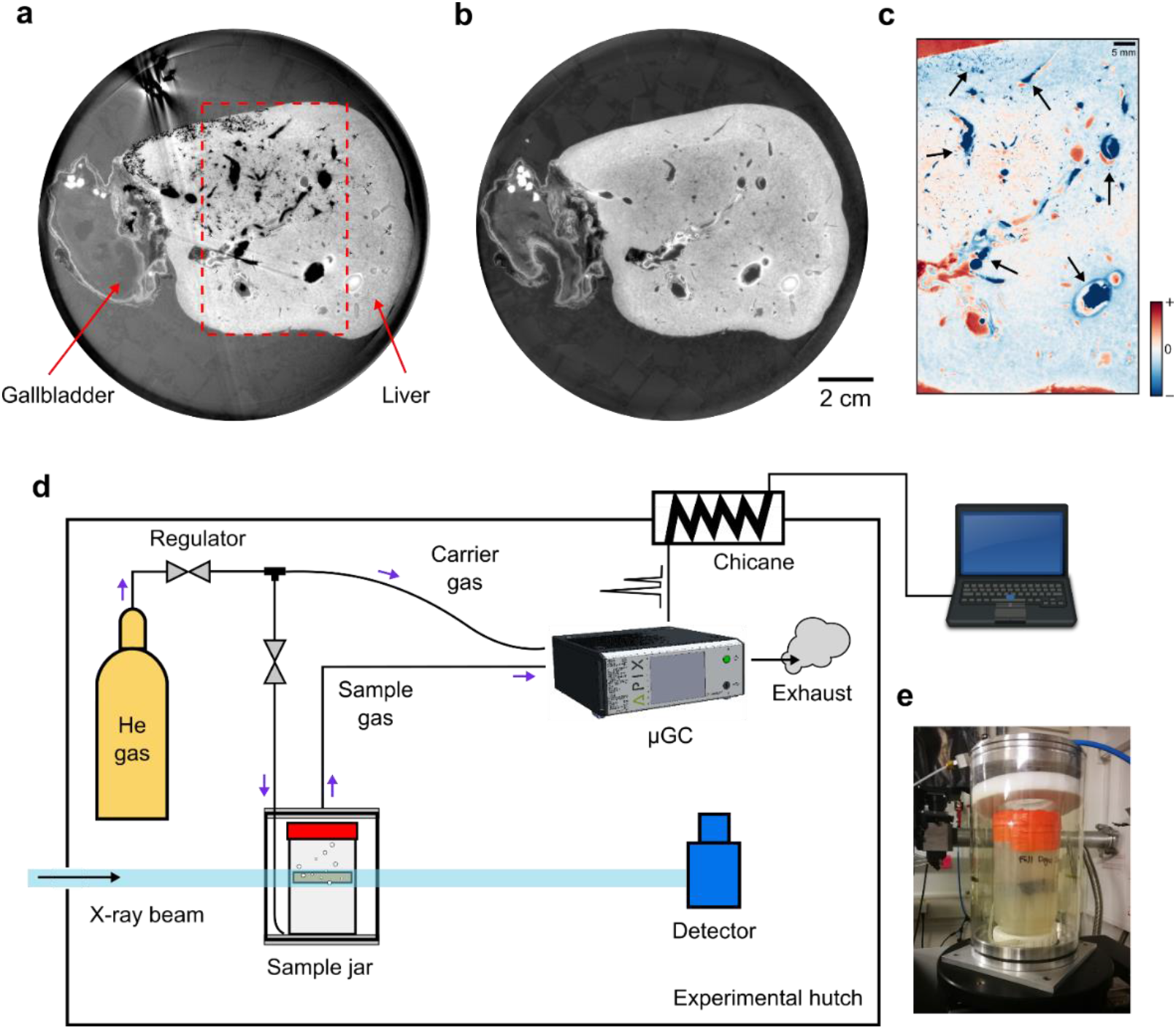
Experimental investigation of X-ray-induced gas bubble formation. **a**, Appearance of bubble formation during X-ray phase-contrast tomography of human liver tissue after a high energy X-ray beam was in the same location for several hours (instead of few minutes) due to scanning equipment malfunction. Compared with **b**, the imaging outcome of the same sample without bubbles (produced by imaging after tissue degassing), the bubbles create huge streaking artifacts in the reconstructed image. Image modified form Brunet et al.[69] **c**, Bubbles revealed (black arrows) by differential intensities between **a** and **b** within the region specified in **a**. The contrast change, such as the negative differential signal within the liver vasculature is primarily caused by bubble formation. **d**, Experimental setup for probing the phenomenon using a micro-gas chromatograph for online detection. **e**, The sample container placed in a sealed external housing.

So far, the conditions and consequences of radiation-induced bubble formation have mostly been investigated in the context of spectroscopy[28] and crystallography[29] at low X-ray energies (< 15 keV), where focused beam geometry in the application contexts greatly intensifies the interaction between X-ray and matter. Moreover, in that energy regime, photochemical processes typically dominate due to the strong X-ray photoelectric effects. For high-energy X-rays (≥ 25 keV)[30], [31], which are suitable for biomedical imaging of large soft tissue samples, little is known about the imaging capacity imposed by the cascading effects from these physical interactions. Especially within the energy range of 60-150 keV, typical for clinical applications[32], high X-ray transmission enables low-dose imaging at a high signal-to-noise ratio using the phase-contrast information of weakly absorbing tissues[15]. Here, X-ray absorption and the subsequent radiation-induced processes are led by Compton scattering compared to lower photon energies, where the photoelectric effect dominates, especially for light elements within soft tissues[23], [33]. For long-term imaging, which can last hours to days, bubble formation has only been briefly mentioned with X-rays in the 20-35 keV range[12], [13]. To our knowledge, no systematic investigation has yet been reported to quantify this phenomenon in the bioimaging context and at high energies very far off-resonance to sample composition.

Understanding the factors influencing X-ray-induced bubble formation is essential for the optimization of sample preparation and imaging protocols to improve the imaging process efficiency and alleviate the failure rate in large-scale projects. To meet this challenge, we used a parallel X-ray beam with typical characteristics for large sample tomography to trigger the bubble formation during in-line phase-contrast cineradiography (time-resolved radiography) of partially ethanol-dehydrated tissue samples. We combined micro-gas chromatography (µGC)[34], [35] with an X-ray imaging beamline to perform online detection of volatile chemical species (such as N_2_, O_2_, H_2_O, and EtOH).

We hypothesized that i) Vacuum degassing during sample preparation would increase the sample resistance to bubble formation, through a reduction in sample dissolved gas concentration ii) More lipid rich tissue would show a lesser reduction in bubble formation due to vacuum degassing, iii) Bubble growth would be constrained by the geometry of the tissue microstructure in which they formed. From the analysis of the two simultaneous time-resolved imaging and µGC measurements, we identified two factors contributing to the bubble formation: i) residual dissolved gas in the samples; ii) X-ray-induced vaporization of the solvents. We also found that in degassed samples bubble nucleation was delayed, that the final bubble load was lower but that these bubbles grew faster initially. The results map out the dose regime for uninterrupted phase-contrast imaging of soft tissues. Moreover, by analysis of global and local bubble dynamics, we identify bubble nucleation sites and distinguish site-specific bubble dynamics pertaining to the geometric constraints in the surrounding space. The collective experimental evidence points to a nonthermal origin of the bubble formation process driven by photoionization of the solvents.

## Results and discussion

In the present study, we monitored the nucleation and growth of gas bubbles in soft tissue samples triggered by polychromatic synchrotron X-ray irradiation centered at 82 keV. Simultaneously, the X-ray beam was also used for phase-contrast cineradiography of the samples accumulated through free-space propagation of the transmitted beam over ∼ 5 m of air in the experimental hutch[6] (see Fig. 1d). In total, two thick slices of human lung and brain tissues, obtained from similar anatomical locations in each organ, were used for this study, without exogenous staining, along with two non-tissue controls filled only with agar-ethanol mixtures (see Materials and methods, SI Fig. 1 and section S1), in each case one of the two samples was vacuum degassed during preparation (see Materials and methods). Bubbles appeared in all tested samples (see SI Fig. 2) after at most ∼ 21 mins of X-ray irradiation. The measured static radiographs and cineradiographs were preprocessed to enhance the contrast of relevant sample composition (agar, tissue, and gas bubbles) by multiple flat-field correction methods (see SI section S2). A compilation of characteristic radiographic frames for a lung sample is shown in Fig. 2 with more shown in SI Fig. 3.

**Figure 2.**
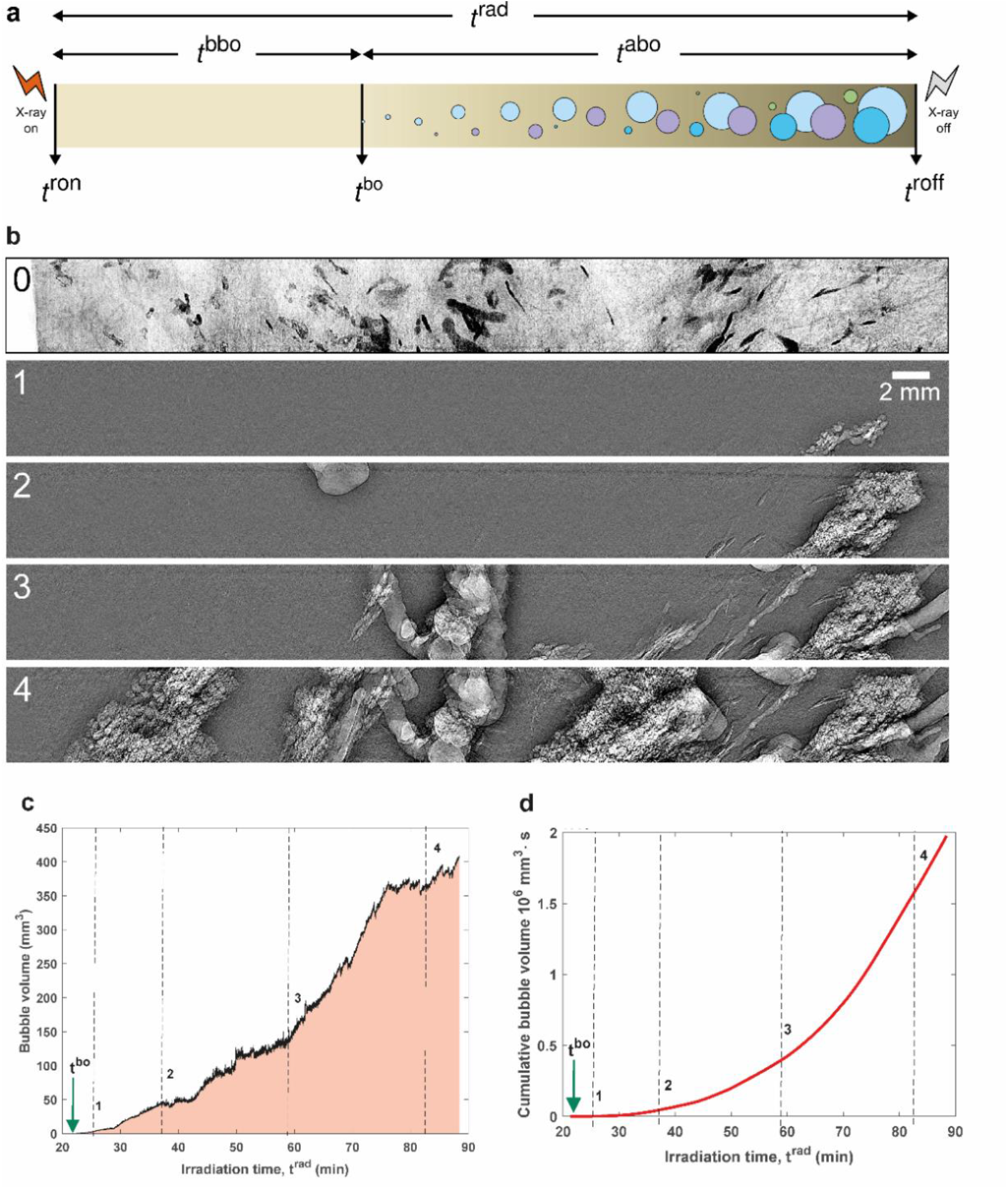
**a**, A schematic illustrating the timescales observed in the experiments. The key time points include radiation on and off times (t^ron^ and t^roff^), and bubble onset time (t^bo^). The time courses include irradiation time (t^rad^), the time before and after bubble onset (t^bbo^ and t^abo^). Bubble that emerged sequentially (using round ones as examples) are indicated with different colors to distinguish one another. The in-line phase-contrast radiographs in **b** each has a field of view of 50 mm (width) by 4 mm (height). Frame 0 shows the lung tissue context in a static radiograph, while frames 1-4 are cineradiographs at different time points after bubble onset. All images in **b** are obtained after context-specific flat-field corrections (see SI section S2). **c**, Quantification of the global bubble evolution using their time-dependent volume within the field of view and **(d)** the cumulative volume. The temporal locations of the frames in **b** are indicated in **c and d** as vertical dashed lines.

To unambiguously describe bubble dynamics, we define four terms, *t*^bo^-(time of bubble onset), *t*^bbo^ (time before bubble onset), *t*^abo^(time after bubble onset), *t*^rad^ (irradiation time), shown in the schematic in Fig 2a. From our observation, bubbles emerge and grow sequentially at multiple sites, (see Fig 2a and Supplementary Videos). An example of bubble formation and evolution is shown in Fig. 2b-d for a human lung tissue sample, where the bubbles first appeared in the alveoli. We quantify the bubble volume growth using the instantaneous time-dependent bubble volume (within the field of view), *V*(*t*) Fig 2c, or the cumulative bubble volume (which includes bubbles which leave the FoV, see Fig. 2d), *V*_*C*_(*t*). They are related according to

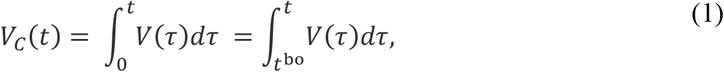

where *t*^bo^ refers to the bubble onset time (see Fig. 2a). We observe that the overall bubble growth exhibits a power-law dependence over time and tends towards saturation in the end. The turning point at around 76 mins since the start of irradiation corresponds with the time point where most of the lung tissue’s air spaces within the field of view of the X-ray beam are filled with gas bubbles. To quantify *t*^bo^, we manually selected the cineradiograph when visible bubbles first appeared within the field of view of the X-ray beam.

### Degassing increases time to nucleation and reduces bubble load for a given X-ray dose

*t*^bbo^ shown in Fig. 3a indicate that the degassing procedure substantially increases the time before bubble nucleation for human brain and lung tissue, as well as for the non-tissue controls containing only the agar-ethanol mixture. The increase of *t*^bbo^ in degassed compared to non-degassed tissues demonstrates quantitatively that vacuum degassing is essential for avoiding bubble formation during prolonged X-ray scans. Over the course of a scans, at any given irradiation time *t*^rad^ the degassed samples consistently have a lower cumulative gas volume (Figure 3c) and the excess gas volume is positive (Figure 3b), there is an exception to this patter for a short time interval in the brain sample. Moreover, the difference between degassed and non-degassed samples is more pronounced (in both nucleation time and difference between highly and non-degassed samples) in the tissue samples derived from the human lung than those from the brain, indicating a relation to their difference in gas solubility and tissue microstructures[36]. These findings support hypotheses (i) and (ii) – suggesting that decreasing the dissolved gas concentration in the sample through vacuum degassing, delays the onset of bubble formation during scanning and that this preparation method is sample composition and microstructure dependent.

**Figure 3.**
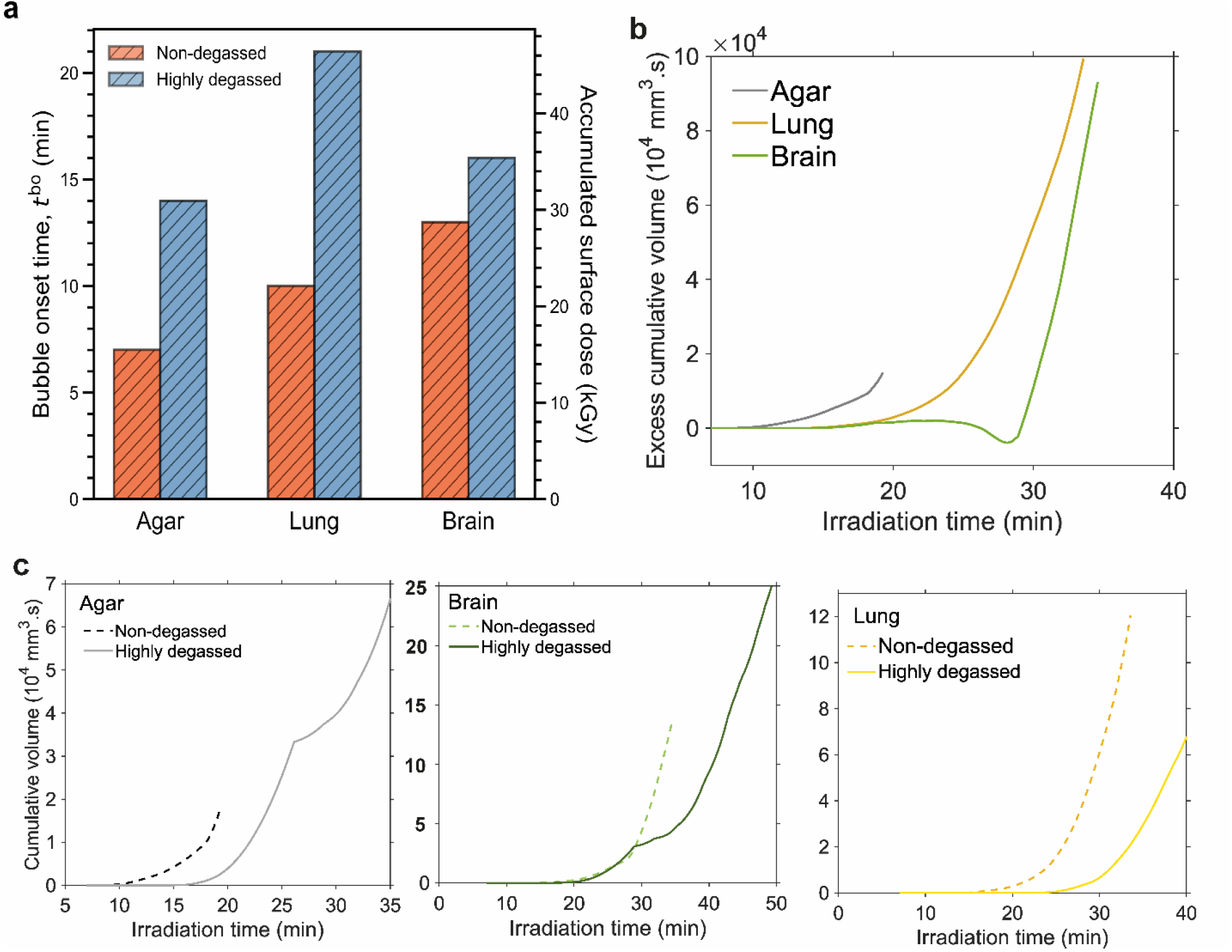
Quantitative characteristics of bubble onset and sample dependence after X-ray irradiation. **a**, Bubble onset time (t^bo^ in Fig. 2a) and X-ray dose deposited into different samples. Compared with non-degassed samples, the degassed samples show markedly delayed bubble onset and therefore higher dose threshold associated with bubble formation. **b**, bubble growth over the entirety of the scan, demonstrating lower cumulative gas volume in the highly degassed samples compared to non-degassed samples. **c**, The difference in cumulative gas volume between non-degassed and highly degassed samples.

### Once formed, bubbles in highly degassed sample initially grow more quickly than non-degassed

For all three pairs of tissue, the bubble time course dynamics can be adjusted for the time of bubble onset to facilitate comparison of the early bubble growth dynamics. After adjusting for the bubble onset times (see Fig. 4a), we observed a faster initial bubble growth rate in highly degassed than non-degassed samples (Fig 4b). To quantify the initial growth trend of X-ray-induced bubbles, we approximate the cumulative bubble volume by a power-law relation,

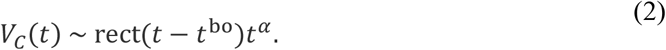

**Figure 4.**
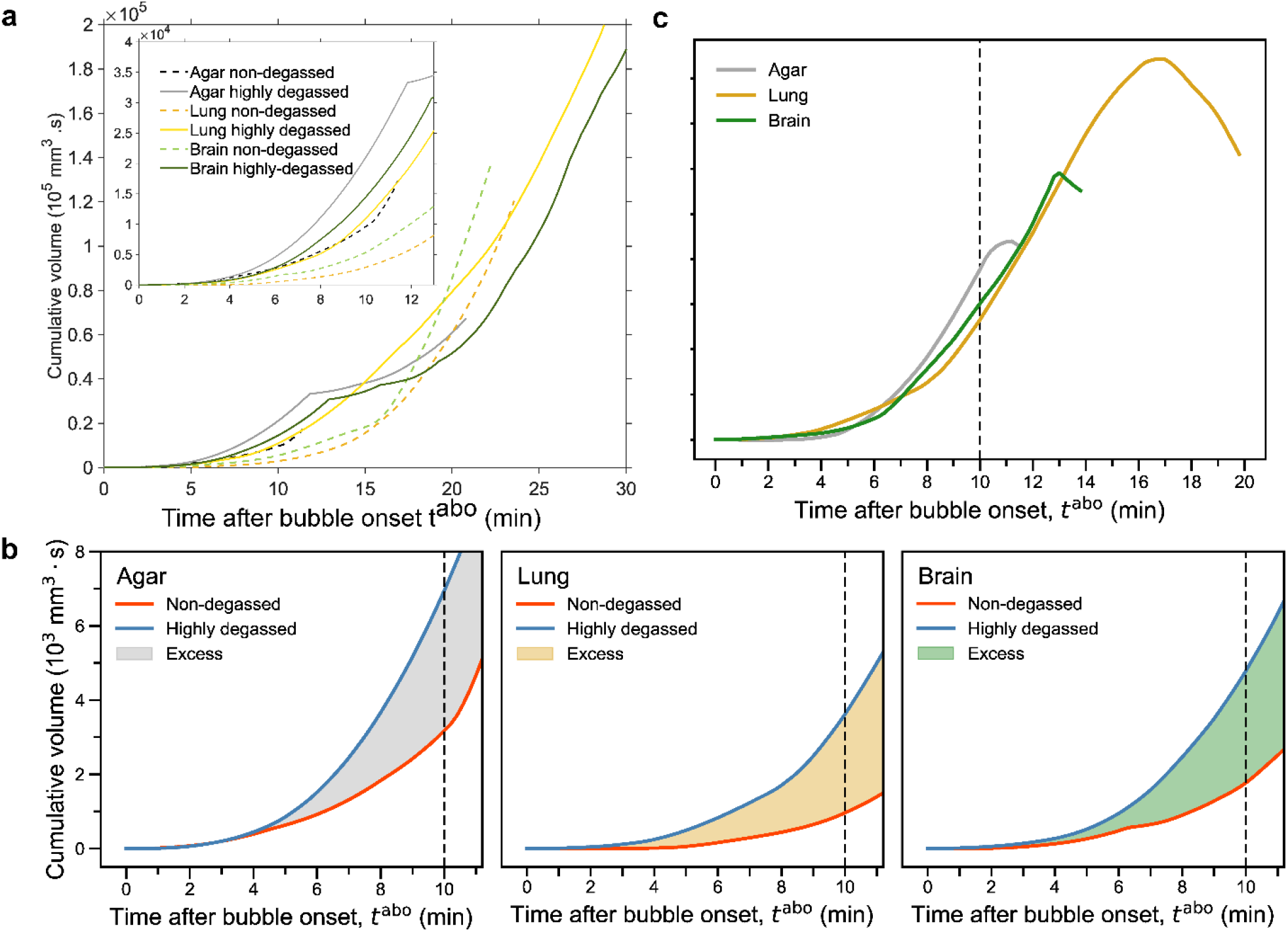
Quantitative characteristics of bubble growth with time adjusted for bubble onset **a**, Bubble cumulative volume for all samples with time courses adjusted for bubble onset time. Inset shows the first 12 mins of bubble growth for all samples. **b**, the initial ∼ 11 mins of bubble growth measured against the time after bubble onset (t^abo^ in Fig. 2a) showing a speed-up for highly degassed samples. **c**, The excess in cumulative bubble volumes for three settings is calculated as a function of t^abo^, showing an increase until saturation, followed by a downward trend. The color choices in **a**-**c** are consistent to display their interrelationship.

Here, rect is a window function, the terms *t*^bo^ represent the bubble onset (bo) time and *α* is the growth exponent. Fitting of Eq. (2) for up to 10 mins after bubble onset yields a consistent *α* ∼ 3.0 for highly degassed samples. Whereas for non-degassed samples, the growth exponent changes by at least 0.4 and is highly dependent on the sample (see SI Table 2). Since dissolved gas modifies the liquid’s thermodynamic landscape[37], the noticeable change in kinetics between highly degassed and non-degassed samples indicates the distinct metastable states the embedding solution was in immediately before bubble onset. In addition, we calculated the difference in terms of cumulative bubble volume as shown in Fig. 4c. For all control pairs, the excess increases until a certain threshold is reached before it drops.

### Bubble growth is geometrically constrained by sample microstructure

When bubbles nucleate in less constrained spaces, such as on the material (tissue and crushed agar) surfaces or in the solution phase of the embedding media (ethanol), they aggregate, coalesce, and often move upwards out of the imaging field of view. These less spatially constrained bubbles tend to be more globular and grow more uniformly. Whereas the bubbles formed within void spaces in the tissue interior such as empty blood vessels or the lung’s air-contacting bronchi are more likely to be trapped *in situ* (bubble entrainment). The entrapped bubbles take on the shape of the surrounding material confining their growth and are generally highly nonspherical. These distinct bubble dynamics all contribute to the overall growth shown in Figs. 2-4. Once the X-ray beam is shut off, the bubbling process keeps developing but gradually subsides as the excess energy dissipates, which can take several minutes in our experiments, as checked by taking static radiographs at later times. The appearances of samples before and after X-ray irradiation are compared in SI Fig. 2, where bubbles are visible around and above the tissue sample up until the container lid.

Understanding the site-specific bubble dynamics can help elucidate the nature of the bubble formation process[38]. To this end, we selected representative bubbles from each sample that were spatially separated from other bubbles within the recorded radiographs (see Fig. 5). For each case, we calculated the bubble circularity (see Materials and methods) and area over time after image segmentation. A circularity of 1 indicates a perfect circle, whereas the closer it approaches 0, the more eccentric the shape becomes. From the shape analysis of the bubble growth, three distinctive scenarios emerge (see Fig. 5): (i) Unrestricted 3D growth of bubbles that nucleated within the gaps of crushed agar, showing a nearly constant circularity over time. The relationship between bubble area (*A*) and the time after bubble onset (*t*^abo^, here referring to the specific bubble’s timescale) is *A* ∼ *t*^abo^, indicating that its radial growth rate 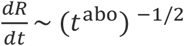, which is consistent with vapor[39] or diffusion-driven growth[38]. (ii) Growth within a large blood vessel in the brain tissue (∼ 100 µm diameter), showing a continuous decrease in circularity that correlates with bidirectional expansion within the tubular vessel, which resembles the behavior of intravascular gas embolism[40]. (iii) Stepped growth of bubbles that indicates the fractal-like expansion within the lung’s peripheral air spaces[41]. The step-like feature in time-dependent circularity and bubble area correlate with the sequential expansion of the bubble into nearby alveoli further and further away from the nucleation site. Gas expansion within neighboring pulmonary alveoli is the fastest among the three scenarios (increasing by ∼ 4×10^6^ µm^3^ in less than 2 minutes as in Fig. 5c) due to their optimized anatomical design that facilitates alveolar gas exchange[42].

**Figure 5.**
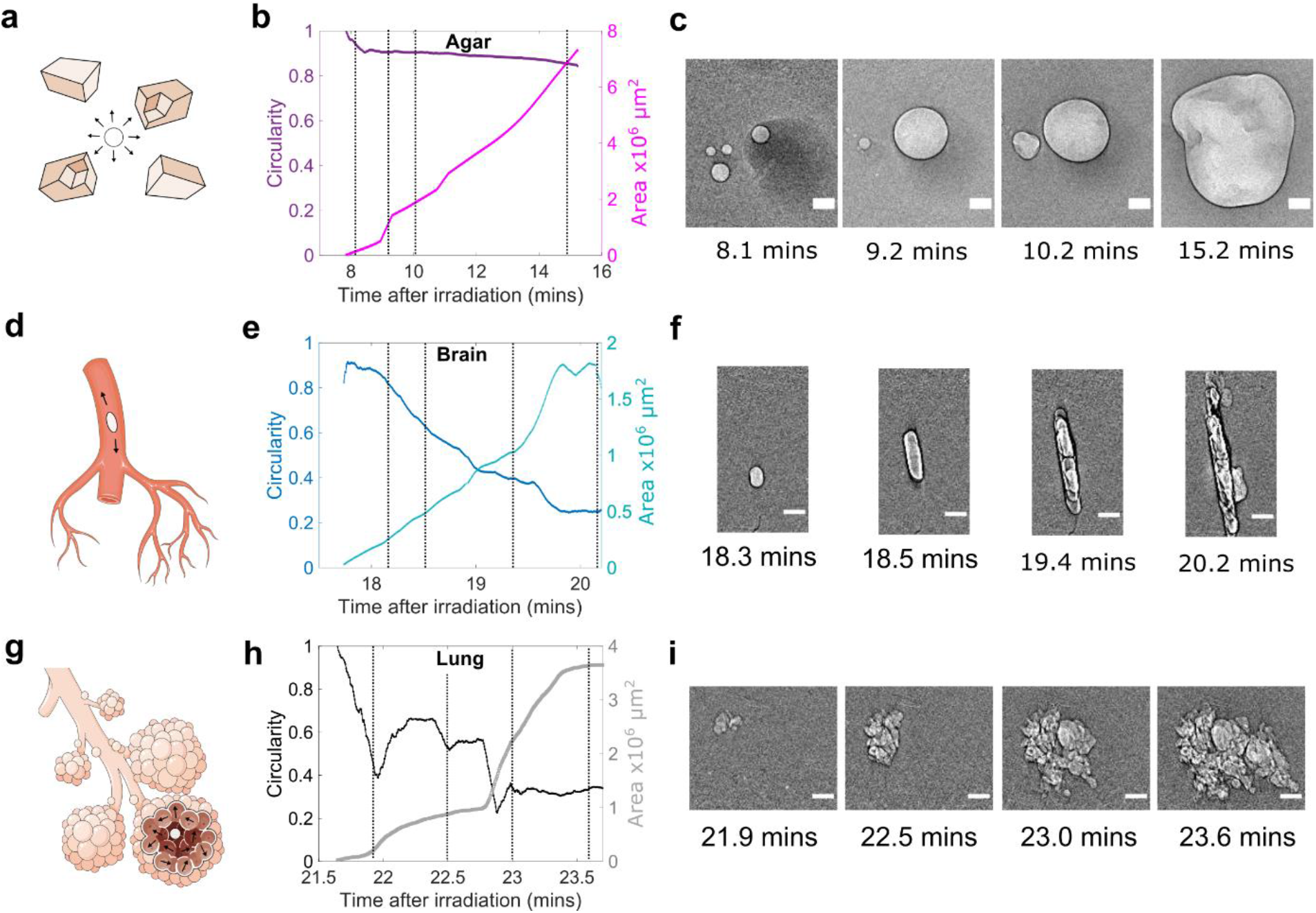
Constrained bubble dynamics in different microscopic void spaces, including **a**-**c**, regions between crushed agar **d**-**f**, within brain vasculature and g-**i**, within alveoli of the lung. **a**,**d**,**g** Schematics showing the local spatial geometry where the bubbles (white or light grey inside in phase contrast) are formed. **b**,**e**,**h** The extracted growth dynamics of the bubbles within these spaces are quantified using time-dependent circularity and bubble area. **c**,**f**,**i**. representative radiographic frames show the morphological change of the specific bubble dynamics. The respective time points are indicated by vertical dashed lines in **b**,**e**,**h**, respectively.

### Gas detection and analysis show changes in relative solvent concentration during bubbling

A brief description of the µGC-based gas detection setup is provided in Materials and methods. The detected gases by the micro-gas chromatograph come from two sources: (i) the dynamical headspace within the plastic housing around the sample container, and (ii) the gases generated within the sample container, including dissolved gas, vapor, and gaseous or volatile end products from potential photoreactions. Chromatographic monitoring provides a way to examine the bubble content as they escape the system on the second to minute timescale. Therefore, the detection scheme naturally filters out short-lived, highly reactive reaction intermediates as well as trace gases since none of them is a main contributor to bubble formation macroscopically. In our case, four detection modules were used in the micro-gas chromatograph to cover different ranges of chemicals (see SI section S4.1). As presented in Fig. 6, the chromatograms show clear signatures of evaporated solvent (EtOH and H_2_O) and gases from ambient air or dissolved gases in tissue (primarily N_2_ and O_2_). Although dissolved O_2_ and N_2_ concentrations differ with tissue composition [36], we have observed that the relative concentrations of N_2_ and O_2_ detected by µGC are essentially unchanged in all cases before and during sustained bubbling under X-ray, as shown in Fig. 4. This contrasts with the behavior of evaporated solvents from µGC. Specifically, EtOH shows an increase in its relative concentration for all samples except the non-degassed agar, whereas it generally decreases for H_2_O. For each sample, the changes took place around the corresponding observed *t*bo, which is largely consistent for all pairs of samples, without a noticeable dependence on sample type. This indicates that the gas from X-ray induced bubbles largely comes from solvent vapor. Moreover, within our detection sensitivity limit, no significant signals of other chemical species have been detected, indicating the relative radiation stability of agar and solvents (see SI section S4.2).

**Figure 6.**
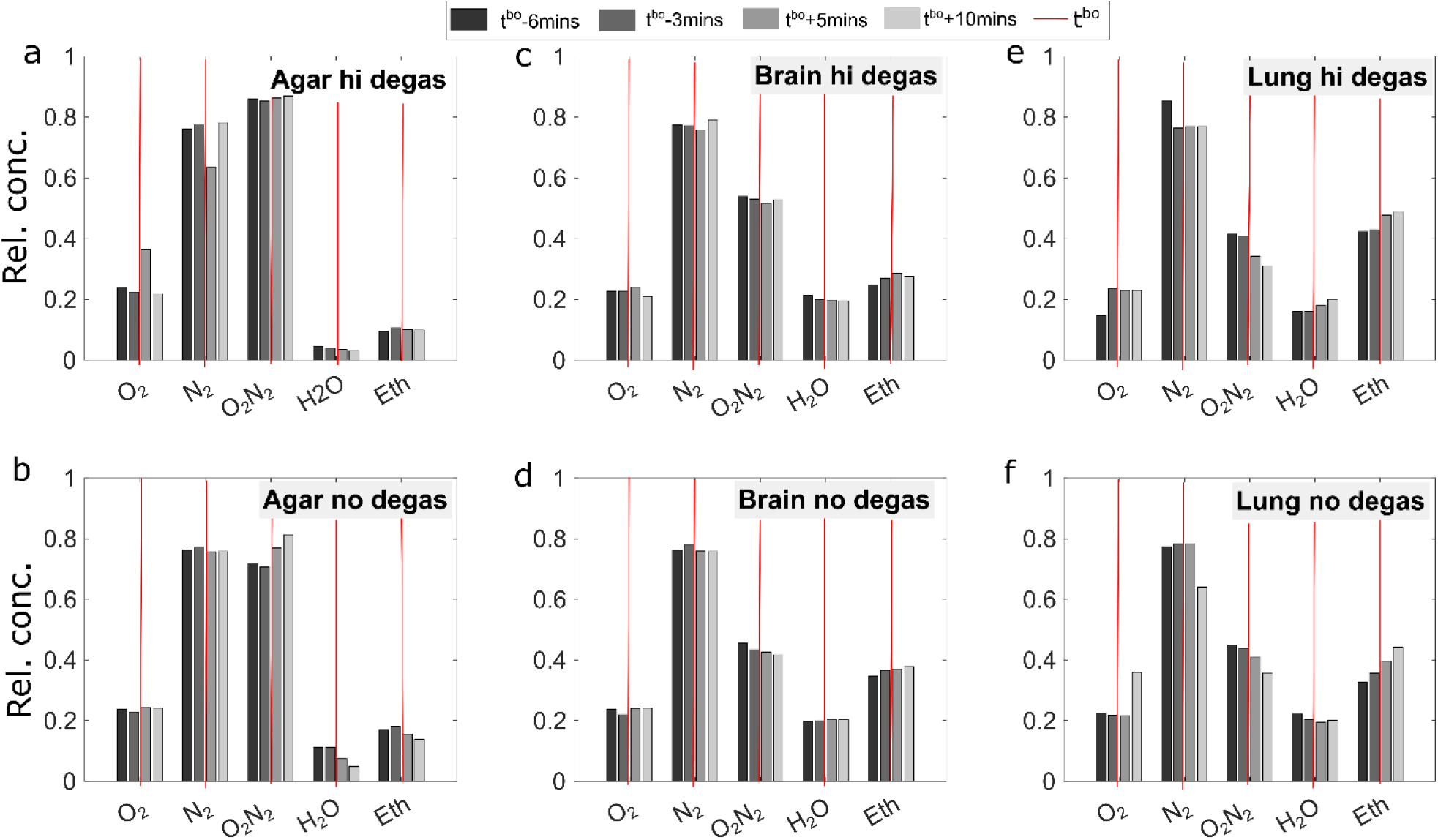
Gas evolution around the bubble onset time (t^bo^) during X-ray irradiation for **a-b**, agar samples, **c**-**d**, human brain tissue samples, and **e**-**f**, human lung tissue samples. Four dominant chemical species detected by the micro-gas chromatograph, including N_2_, O_2_, EtOH, and H_2_O, are shown here in their relative concentrations (rel. concentr) at 6 mins, 3 mins before bubble onset and 5 mins, 10 mins after it. The relative concentrations are separated into two sets corresponding to the two modules: MS5A measured O_2_ and N_2_ concentrations, while PDMS10 measured EtOH, H2O, and 0_2_+N_2_. For each type of sample, the corresponding t^bo^ is drawn in the figures with vertical red lines. The general behavior before and after bubble onset is that the relative concentrations of dissolved gases (N_2_ and O_2_) remain constant, while the solvent vapor (EtOH and H_2_O) shows noticeable changes.

### Mechanisms driving bubble nucleation and growth

In addition to our initial hypotheses regarding sample preparation and tissue composition, our study enables the investigation of the mechanisms that underlie bubble formation and growth in the high energy X-ray regime. Our quantitative observations for our experimental conditions led us to consider a few plausible underlying mechanisms, which can come from chemistry and heating (thermal or nonthermal). The mechanisms we considered were (i) accumulated photochemical effects, (ii) dose-dependent solvent heating effects, (iii) phase transition in the gel embedding. Regarding scenario (i), the most likely reactions are the radiolysis of EtOH and H_2_O, which produce gases (primary products H_2_, CH_4_, secondary product O_2_, etc), radical ions and oxidized solvent as end products[29], [43]–[45]. Since the high-energy X-rays used for imaging are very far off resonance to sample composition, we expect the gas contribution from direct water and ethanol splitting with X-rays[46] to be low, albeit nonvanishing. This stems from the two-order-of-magnitude reduction of photoelectric effect cross-section between X-rays in the energy range 80-100 keV and 10-30 keV for light elements which constitute soft tissues[33]. In fact, an earlier experiment on ice showed that X-ray-induced H_2_O decomposition is undetectable with X-ray energies above 30 keV[46]. In our case, the reasoning is supported by the µGC results, which identify X-ray-induced vaporization[47] as a primary source of gas escape.

To evaluate (ii) and (iii), we performed dosimetry measurements with an ionization chamber dosimeter and numerical estimation (see SI section S3). With a measured dose rate of 36.8 Gy/s, we found that the accumulated surface dose on the sample range between 15.5 kGy (non-degassed Agar) and 46.4 kGy (highly degassed Lung) just before bubble onset (see Fig. 3a and SI Table 3). Assuming no heat exchange, the thermal load imparted to the sample generally creates a temperature increase within the X-ray field of view of up to about 11 °C, if assuming only H_2_O, or 19 °C, if assuming only EtOH. Although detailed dosimetry calibrations are beyond the scope of this work, our estimation provides a sensible upper bound on these values. Therefore, the macroscopic metastable state at bubble onset differs from the two prominent examples of thermally-induced bubble formation involving electromagnetic radiation: (a) resonant heating which produces plasmonic microbubbles from water vaporization observed in solvated gold nanoparticles[48]; (b) the bubble-chamber phenomenon[49], [50], which requires a superheated liquid medium to be maintained between its boiling and critical temperature for the effect to take place. For 70% EtOH, the boiling point is over 78 °C at ambient pressure[51], which is inconsistent with our experimental condition. However, nonthermal processes through rapid photoionization can produce heating effects through disruption of the coordination environment in the solvation shell[52] or lowering of the evaporation enthalpy[47], [53]. In our case, it is likely that the exsolution of dissolved gas in tissue due to a moderate temperature change or trace gas generated through photochemical reactions seeded the bubble formation in the solvent, which is further promoted by a photoionization-disrupted local solvation environment to rupture the bulk liquid.

For scenario (iii), the working hypothesis is that the agar may undergo thermoreversible sol-gel transition due to X-ray-induced heating, resulting in the release of entrapped gas. This behavior has been observed at much lower X-ray energies (∼ 30 keV), resulting in melting of the embedding agar gel during imaging[13]. However, in our case, the crushed agar has been degassed after gelling and before its use in sample preparation. At the considerably higher X-ray energy we used for radiography, no change in the consistency of the agar gel was observed from before the X-ray is switched on and immediately after the beam is shut off. Moreover, due to the large thermal hysteresis of agar, its melting temperature is much higher than the gel-setting temperature, therefore the very moderate temperature change we estimated before *t*bo would unlikely initiate melting.

## Discussion

Our estimation of the bubble onset and volume estimation are only accounting for those within the field of view of the X-ray beam, hence we excluding the bubbles generated from scattered X-rays. Nevertheless, we observe that X-ray-induced bubble formation appears to have a dose threshold in the 10^4^ kGy range for in-line radiography. Vacuum degassing delays the bubble onset time, and reduced bubble load over timescale of a typical scan. The degassing process also leads to faster bubble growth than non-degassed samples at this delayed time. These results, on the one hand, complement existing discussions of the dose-resolution trade-off in X-ray nanoimaging of single cells and thin tissues and in macromolecular crystallography[3], [11], where samples (at only up to millimeters in size) are typically kept in a frozen-hydrated or freeze-dried state. On the other hand, they relate to the emerging field of X-ray virtual histology[6], [15]–[17], where the balance of sample integrity and imaging throughput requires consideration of radiation-matter interaction across various scales determined by the biological questions.

Since bubbles can cause local deformation which may lead to damage in soft materials including tissues[26], [27] from the pressure exerted on them during cavitation, collapse or subsequent bubble entrainment, our results provide the empirical foundation for developing precautionary measures, such as tissue degassing, pressurization, temperature stabilization, to mitigate or postpone bubble formation, thereby reducing the measurement failure rate. The methods described here may also be used to further investigate microbubble formation and growth phenomena in various tissues relevant for clinical models. The examples include air embolism[54], [55] from positive pressure ventilation or after scuba diving accidents, where alveolar gas ruptures into neighboring capillaries due to an increase in gas volume[56], and decompression sickness, where the mechanism and progression of off-gassing from tissue and vasculature remains unproven[57]. In both cases, constructing microscopically accurate physiological simulations require empirical parameters that are often hard to obtain in parenchymal tissues by conventional means.

Beyond the context of bioimaging, spontaneous bubble formation during X-ray experiments has also been briefly reported in X-ray spectroscopy[28] and *operando* imaging of electrodeposition[19] and batteries[58] involving liquid electrolytes. In these scenarios, bubbles likewise produce undesired and sometimes deleterious effects for the precise characterization and stable operation of devices. However, because of the complexity of the liquid environment and gases produced during electrochemical operations[59], it remains exceedingly difficult to separate the sources of bubbles. Our observations in soft biological tissues and gels reaffirm the proposition that X-rays may not be treated solely as a passive probe under intense or prolonged irradiation[23], although isolating bubble sources will require subsequent studies in simplified model systems. The set of control experiments reported here demonstrate that bubble formation remains an issue at high X-ray energies, when sufficient dose is deposited into the sample and its liquid/gel environment, regardless of the initial condition.

## Conclusion

As we continue to push X-ray imaging to higher resolutions on ever larger samples[60], [61], probing the dose limits of bubble formation and developing mitigation methods, either in instrumentation, measurement protocol or data processing, becomes increasingly important. Here, we have combined *operando* gas chromatography with in-line X-ray phase-contrast cineradiography to investigate bubble formation under high-energy X-ray irradiation. We observe the quantitative growth characteristics of X-ray-induced bubble formation from the microscopic to the macroscopic level and have shown how degassing can substantially delay bubble nucleation and thereby lengthen the duration of uninterrupted imaging.

Our current identification of the stable, long-timescale gas products from X-ray irradiation invites further investigation into the corresponding short-timescale mechanism down to the molecular level, preferably in simplified systems where multiple probes may be simultaneously used. Collectively, they will facilitate model-building for radiation-matter interaction and bubble dynamics in complex media from first principles[62], or using continuum models coupled with local geometry[63]. Furthermore, the study illustrates the potential of X-rays as simultaneously a probe and an initiator of bubble dynamics, future research should investigate more efficient degassing methods and quantify the dissolved gas concentration in tissue, as well as continue the efforts to reduce the X-ray dose while keeping or even increasing the data quality. The high sensitivity and resolution of modern phase-contrast X-ray imaging offers an appropriate tool for soft materials rheological characterization[26], [64] using bubbles as the local probe, which may be carried out in opaque media that are traditionally not amenable to study with optical imaging means. In these contexts, our results motivate the continued development of high-speed and multiscale tomography[65] and experimental designs to sample ensembles of diverse bubbles in 3D[66] with a large field of view. When combined with energy-dependent studies and *in situ* thermometry[67], our approach will uncover the progression and characteristics of radiation-matter interaction on the intermediate spatiotemporal scale.

## Materials and methods

### Sample preparation

The lung and the brain used for the experiment obtained from two organ donors registered at the Laboratoire d’Anatomie Des Alpes Françaises (LADAF) in Grenoble, France. Dissections were performed respecting current French legislation for body donation. Body donation was based on free consent by the donors antemortem. All dissections respected the memory of the deceased. The post-mortem study was conducted according to QUACS (Quality Appraisal for Cadaveric Studies) scale recommendations[68]. The postmortem processing of organs (human brain, lung, and liver) follows the previously discussed procedure involving formalin fixation and ethanol dehydration[6], [25], [69]. The human liver sample was imaged twice with HiP-CT, an X-ray phase-contrast imaging technique compatible with large soft tissue samples[6]. After the initial bubble formation due to a beam crashed with the beam on for 4 hours (instead of a normal scanning time of 10 minutes), the sample was re-degassed and re-imaged for the second round. The human brain and lung samples were cut into ∼ 20 mm-thick slices with a diameter of ∼ 70 mm (see SI Fig. 1). The slices fit laterally into the cylindrical plastic containers such that the section plane is parallel with the lid to simulate the imaging geometry in tomography[6] and to facilitate gas escape as they are formed on the tissue interior and surfaces. The tissue slices were immersed in 70% ethanol solution in water and immobilized with crushed agar[6], [25], [69]. The non-tissue controls contain only the agar-ethanol mixture. Details on the sample preparation and mounting are provided in the SI section S1.

### In-line phase-contrast cineradiography

X-ray phase-contrast radiography and cineradiography experiments were carried out at the ESRF BM05 beamline using a polychromatic source. The X-ray beam dimensions were 51.2 × 5.2 mm and was attenuated by 40 mm SiO2 and 1.53 mm aluminum for an average energy of 74 keV. The radiographs were directly recorded with the parallel beam configuration shown in Fig. 1c using a 2 mm-thin LuAG: Ce scintillator (Lutecium-Aluminum Garnet doped with Cerium, custom-made by Crytur, Czechia) and an sCMOS light sensor (PCO edge 4.2 Camera Link HS, PCO Imaging, Germany). The measurement configuration has a propagation distance of 3475 mm, resulting in an isotropic pixel size of 25 µm for all radiographs. For each sample container, we started by locating the sample position with static phase-contrast radiographs. To trace the onset and evolution of X-ray-induced bubbles, cineradiographs were recorded with a data acquisition rate of 3 images per second for at least ∼ 10 minutes after the first bubble appeared within the field of view.

### Operando gas chromatography

In the experiments, we used a portable commercial micro-gas chromatograph (ChromPix2, APIX Analytics, France) along with the data acquisition software DataApex Clarity v8.5. Four detection modules (MS5A, PDMS5, PDMS10, PPU), each containing a separate column as the stationary phase, were used to cover a wide range of detectable molecular species. We recovered the relative concentrations of EtOH and H_2_O from the µGC module PDMS10, and those for N_2_ and O_2_ from the module MS5A. To improve gas detection, a reservoir of He gas was used as the carrier gas to continuously flush the volume within the cylindrical housing (see Fig. 1d and SI Fig. 2a), which functions as a dynamical headspace, to inject the escaped gas into the chromatograph. A pump is attached to the exhaust of the µGC to facilitate the gas extraction. During operando measurements, each round of gas elution through the four detection modules requires about 2.4 mins.

### Computational analysis

Multiple flat-field images from static radiographs and cineradiographs have been used to quantify the bubble volume. Quantification of the bubble volume is possible after calculating and subtracting the absorption profile of the agar and tissue from the overall profile. The volume calculation used the direct inversion of the Beer-Lambert law (see SI section S2). The bubble circularity in constrained environments is calculated using *4πA/C*^2^, where *S* and *C* are, respectively, the area and perimeter of the specific bubble obtained from image segmentation.

## Data, Materials, and Software Availability

The complete imaging data generated and analyzed in the current study are available from the corresponding authors upon request.

## Acknowledgements

We thank P. Masson and A. Bellier at LADAF in Grenoble, France, for collecting organs from the human donors. We thank Philippe Sabot and Régis Barattin from APIX Analytics (Grenoble, France) for their help with the gas chromatography. We thank the scientists sharing their knowledge and past experiences on the bubbling phenomena they encountered during synchrotron beamtimes. These people include A. Mittone at ALBA Synchrotron (Barcelona, Spain), P. P. Paul, M. di Michiel and M. Krisch at ESRF, B. Bay at Oregon State University (Corvallis, OR, USA), C. Disney at University College London. R.P.X. thanks R. Santra at the Center for Free-Electron Laser Science (Hamburg, Germany) and S. Carbajo at SLAC National Accelerator Laboratory (Palo Alto, CA, USA) for helpful discussions. In addition, we thank S. E. Verleden at the Antwerp Surgical Training, Anatomy and Research Center (ASTARC) in Belgium for helpful discussion. We thank H. Müller and P. Lloria at ESRF for lending of equipment. This project has been made possible in part by grant number CZIF2021-006424 from the Chan Zuckerberg Initiative Foundation, grant 2020-225394 from the Chan Zuckerberg Initiative DAF, an advised fund of Silicon Valley Community Foundation, The ESRF - funding proposal md1252, and the Royal Academy of Engineering (PDL - CiET1819/10) and the MRC (MR/R025673/1).

