## Supplementary information for "A closer look at high-energy X-ray-induced bubble formation during soft tissue imaging"

### Supplementary Information for
### A closer look at X-ray-induced bubble formation in soft tissue imaging

R. Patrick Xian[1,4,†,*], Joseph Brunet[1,4,†], Yuze Huang[1], Willi L. Wagner[2,3], Peter D. Lee[1,*], Paul Tafforeau[4,*], Claire L. Walsh[1]

**[1]Department of Mechanical Engineering, University College London, London, UK.**

**[2]Department of Diagnostic and Interventional Radiology, University Hospital Heidelberg, Heidelberg, Germany.**

**[3]Translational Lung Research Centre Heidelberg (TLRC), German Lung Research Centre (DZL), Heidelberg, Germany.**

**[4]European Synchrotron Radiation Facility, Grenoble, France.**

**[†]These authors contributed equally to the work.**

### S1. Samples before and after X-ray radiography

#### S1.1. Sample preparation and mounting

The sample preparation procedure follows the prior description for whole organ imaging(1), which we summarize in brief here: The harvested organs from two human donors were fixed with 4% formalin solution followed by ethanol (odorless bioethanol, Cheminol, France) with increasing concentration until 70% in water(1, 2). Before sectioning into thick slices, the samples were gently dried with blotting papers. The brain slices were taken from the cerebrum, while the lung slices were taken from the upper lobe of the right lung. Photographs of the sliced tissue samples prepared for the experiments are shown in Supplementary Fig. 1a-b.

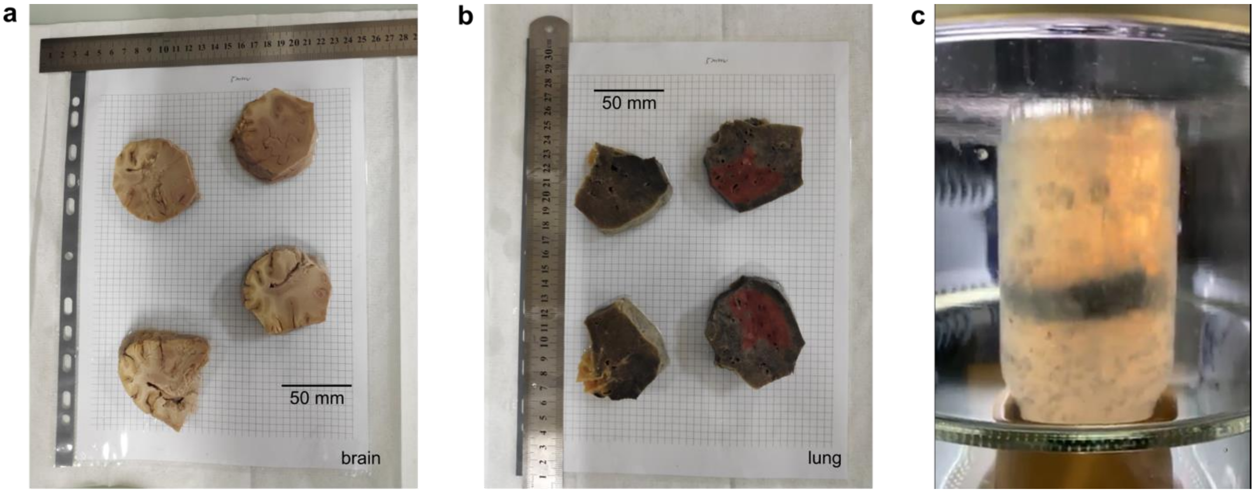

**Supplementary Fig. 1** Thick tissue sample slices of parts of **a**, a brain and **b**, a lung from human donors used in the experiments. The tissues were fixed in 4% formalin and increasing concentrations of ethanol (up to 70%) before slicing. The grid size beneath the sample is 5 mm along each side. **c**, A slice of lung tissue mounted in the plastic container (immobilized by crushed agar in 70% ethanol solution) is being degassed in a glass dryer attached to a pump from the top.

Of the six sample containers, four of them were half-filled with crushed, degassed agar gel (made from powder purchased from Nat-Ali, France, see Ref. (1) for details), then the sample was placed in the container, before filling to the top with the mixture of crushed agar and 70% ethanol. Then, one sample of each tissue type (brain and lung) and a control sample with undegassed agar gel were put aside and labeled as "non-degassed" samples. The remaining three containers were degassed using a diaphragm pump (Vacuubrand, MV2, 1.9 m3 h−1) in a glass dryer (see Supplementary Fig. 1c). While degassing, the volume loss due to the removal of dissolved gas and solvent evaporation was replenished with proportional aliquots of crushed agar-ethanol mixture from a reservoir. Upon finishing, these samples were referred to as "highly degassed". Degassing was terminated using both visual assessment and quantitative information. Visually, good degassing leaves behind only a small number of bubbles in the container and no centimeter-sized bubbles are trapped. Besides, we timed the bubble onset from the start of pumping to quantitatively gauge the quality of degassing. Initially, bubbles appear within about 20 s, while at the end of degassing, the bubbles appear only after 90 s. Finally, after degassing, the sample containers were sealed with parafilm and duct tape, and a small hole (~ 1 mm in diameter) was punctured on the lid to allow gas to escape during irradiation. Until that, this hole was covered with leak tight tape to ensure that no gas from the air could re-dissolve in the samples. After mounting, the samples were kept at 5°C until 12 hrs before radiography measurements, when they were stored at room temperature.

##### S1.2. Samples under and after X-ray irradiation

As shown in Supplementary Fig. 2a, the sample jar is fixed onto the tomograph at the X-ray beamline (ESRF BM05) using a custom-built plastic housing sealed at the top and the bottom with O-rings. The

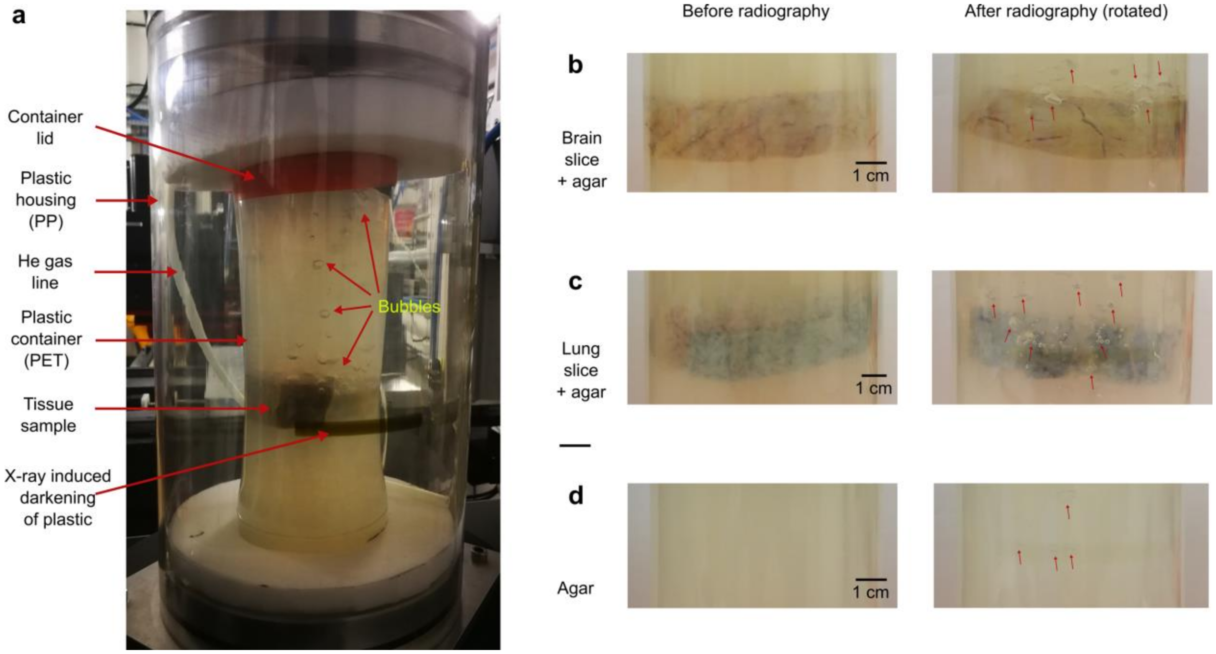

**Supplementary Fig. 2** General appearance of gas bubbles within the experimental setting. **a**, After parallel-beam radiography at the ESRF BM05 synchrotron beamline with polychromatic X-rays (centered around 82 keV), the sample container shows bubbles trapped within the gaps between crushed agar, while the plastic housing show a pronounced darkening mark from X-ray irradiation with the exact shape of the beam. The before and after images of the samples in the container are shown for **b**, a brain slice, **c**, a lung slice and **d**, for the control container with only the embedding medium (crushed agar and 70% ethanol in water). The red arrows in **b**-**d** point towards regions with gas bubbles for visual guidance. The main pictures in **b**-**d** were taken with a phone camera under normal indoor lighting, while the zoom-in pictures of bubbles are contrast-adjusted to enhance the bubble boundaries.

photograph also shows the typical appearance of the sample container and the external plastic housing immediately after X-ray radiography: The dark rectangular mark is a result of color center formation in the plastic upon high X-ray irradiation(3), which is pronounced in PP (material used for the plastic housing), but negligible in PET (material for the sample container). The mark is largely reversible afterwards through annealing, UV light or blue light irradiation, and will tend to fade with time. No evident change in the plastic properties has been noticed so far even after dozen cycles of irradiations. It labels the synchrotron beam location where bubbles are often concentrated: Some bubbles are found at various locations (mostly) above the sample up to the container lid. Supplementary Fig. 2a provides a snapshot of the bubble dynamics.

A second similar mark is visible on the other side of the plastic mounting tube where the beam goes out of the sample. This second mark is less pronounced due to the absorption of the X-ray beam in the sample.

Using optical photographs taken with a phone camera, we compare the appearance of bubble distribution in different tissue samples and the embedding medium before and after X-ray radiography in Supplementary Fig. 2b-d. Although bubbles are present in all cases, they are more pronounced when the tissue slices are present, due to increased chances of bubble formation and entrainment within the tissue interior, such as the vasculature, airways, ventricles as well as complex interfaces. Moreover, the color contrast between the tissue and the embedding and the turbidity of the embedding from light scattering in agar(4) also contribute to the visibility of bubbles in the photos. Nevertheless, their stable yet highly nonspherical shapes, likely stabilized by geometric confinement, become more visible after contrast adjustment (see zoomed-in regions in Supplementary Fig. 2b-d).

The bubble distributions are better visualized in X-ray radiographs as shown in Supplementary Fig. 3, after removing tissue contribution in the signal (see SI section S2). Similar to Main Text Fig. 5, we can use the shape to directly distinguish between bubbles stemming from different nucleation sites. Bubbles with smooth, globular features were found in all three types of samples, which come from the surrounding

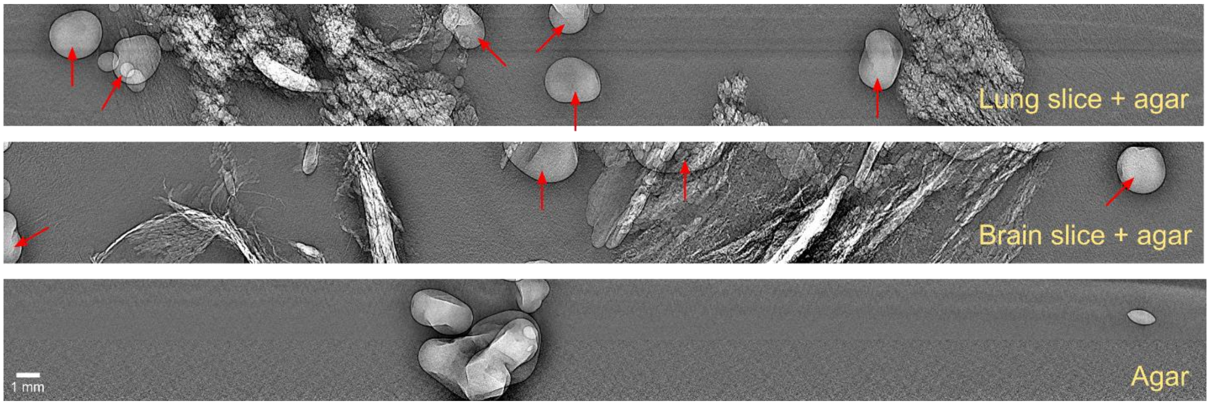

**Supplementary Fig. 3** Comparison of bubble morphologies in phase-contrast radiographs of lung slice (top), brain slice (middle) and in the embedding medium of agar and 70% ethanol mixture in water (bottom). The bubbles nucleated in gaps between crushed agar are indicated with red arrows in the top and middle images. The scale bar of all radiographs is the same as in the case with only agar.

embedding. The bubbles trapped in vasculature has an elongated fiber-like shape, while the bubbles which originated in the lung alveoli have a rugged shape. The formation of these site-specific bubbles is recognizable in the corresponding Supplementary Movies.

#### S2. Radiographic image processing and quantification

For each frame in cineradiography, the transmission signal ($I_{\mathrm{T}}$) registered by the detector has contributions from multiple components. We distinguish two temporal regimes relevant for the data analysis: the time before bubble onset (bbo) and the time after bubble onset (abo), which have been introduced in Fig. 2a in the Main Text. In these two regimes, the transmission signal, including both absorption and phase contrast, is determined by the following relations, respectively.

$$I_{\mathrm{T}}^{\mathrm{bbo}} = I_{\mathrm{dark}} + I_{\mathrm{medium}}, \tag{3}$$

$$I_{\mathrm{T}}^{\mathrm{abo}} = I_{\mathrm{dark}} + I_{\mathrm{medium}}(t^{\mathrm{rad}}). \tag{4}$$

We assume the bubble dynamics create an irradiation time ($t^{\mathrm{rad}}$) dependence of the transmission signal's intensity contribution through attenuating media (i.e. air and sample) along the beam path ($I_{\mathrm{medium}}$), while the contributions from other components are largely independent of $t^{\mathrm{rad}}$. To access the signal contribution from the bubbles, the dark counts of the detector ($I_{\mathrm{dark}}$) were removed directly by subtraction, other contributions were suppressed or removed using multiple flat-field corrections (FFCs)(6, 7) as explained next.

##### S2.1. Multiple flat fields for phase-contrast cineradiography

FFC is traditionally used for compensating the non-uniformity of illumination(8, 9). In X-ray imaging, where transmission is typically measured, FFC is also used for the removal of ring artifacts(6, 7). Here, we use it to isolate the bubble contribution to the signal by treating other intensity contributions in Eq. (3)-(4) as the *de facto* (stationary) background. The usefulness of FFCs is supported by the multiplicative form of the Beer-Lambert law, which we illustrate with a 1D example below.

$$I_{\mathrm{medium}} = I_0 \exp\left(-\int \sum_i \mu_i \, dl\right) = I_0 \prod_i A_i, \tag{5}$$

$$A_i = \exp\left(-\int \mu_i \, dl\right). \tag{6}$$

The component-wise attenuation factors ($A_i$) relate to the linear attenuation coefficient (LAC) of each component ($\mu_i$) according to Eq. (6). The integration is over the beam propagation direction in 1D for this example. After dark count subtraction, the measured X-ray transmission in Eqs. (3)-(4) as may be expressed as a product of different attenuation factors,

$$I_{\mathrm{T}}^{\mathrm{bbo}} - I_{\mathrm{dark}} \simeq A_{\mathrm{air}}A_{\mathrm{agar}}A_{\mathrm{tissue}}, \quad (7)$$

$$I_{\mathrm{T}}^{\mathrm{abo}} - I_{\mathrm{dark}} \simeq A_{\mathrm{air}}A_{\mathrm{agar}}A_{\mathrm{tissue}}A_{\mathrm{bubble}}(t^{\mathrm{rad}}). \quad (8)$$

Although Eqs. (7)-(8) hold exactly only for monochromatic radiation, in the setting of polychromatic X-ray as is used for our experiment, they can still maintain a multiplicative form upon generalization(10). We leave detailed derivation on the complex time-dependent scenario for future work.

**Supplementary Table 1** Flat-field corrections used for resolving different contexts.

| **Type** | **Characteristic** | **Term symbol** | $\boldsymbol{I_{\mathrm{object}}}$ | $\boldsymbol{I_{\mathrm{ref}}}$ | **Usage** |
|---|---|---|---|---|---|
| I | static | $I_{\mathrm{ff,stat}}^{\mathrm{agar,air}}$ | Static radiograph through agar | Static radiograph through air | For quantifying agar absorption profile |
| II | static | $I_{\mathrm{ff,stat}}^{\mathrm{tissue,air}}$ | Static radiograph through tissue | Static radiograph through air | For quantifying tissue absorption profile |
| III | static | $I_{\mathrm{ff,stat}}^{\mathrm{tissue,agar}}$ | Static radiograph through tissue | Static radiograph through agar | For locating sample within the field of view |
| IV | dynamic | $I_{\mathrm{ff,dyn}}^{\mathrm{tissue,air}}$ | Cineradiograph through tissue | Static radiograph through tissue | For quantifying bubble volume |
| V | dynamic | $I_{\mathrm{ff,dyn}}^{\mathrm{tissue,tissue\_bbo}}$ | Cineradiograph through tissue | Cineradiograph through tissue before bubbling | For making movies of bubble dynamics |

The general formulation of FFC may be written as(7, 11, 12)

$$I_{\mathrm{ff}}^{\mathrm{object,ref}} = \frac{I_{\mathrm{object}} - I_{\mathrm{dark}}}{I_{\mathrm{ref}} - I_{\mathrm{dark}}}, \quad (9)$$

where $I_{\mathrm{object}}$ and $I_{\mathrm{ref}}$ are the object-included and reference (i.e. object-excluded) images, respectively. The reference image is commonly referred to as the flat field. In practice, a judicious choice of FFC in X-ray phase-contrast imaging of soft tissues greatly improves the image contrast and enables the visualization of samples exhibiting very low density variations(1, 11). In the series of phase-contrast cineradiography experiments presented in this work, we acquired flat-field radiographs for transmission through the air and the agar embedding at a different location within the same sample container (see Supplementary Fig. 2a). We vary the $I_{\mathrm{object}}$ and $I_{\mathrm{ref}}$ in Eq. (9) for suppression and removal of signal components other than bubbles. In total, six different types of FFCs have been used for the cineradiography experiment and bubble quantification. The details including their respective usage are listed in Supplementary Table 1 and further explained afterwards.

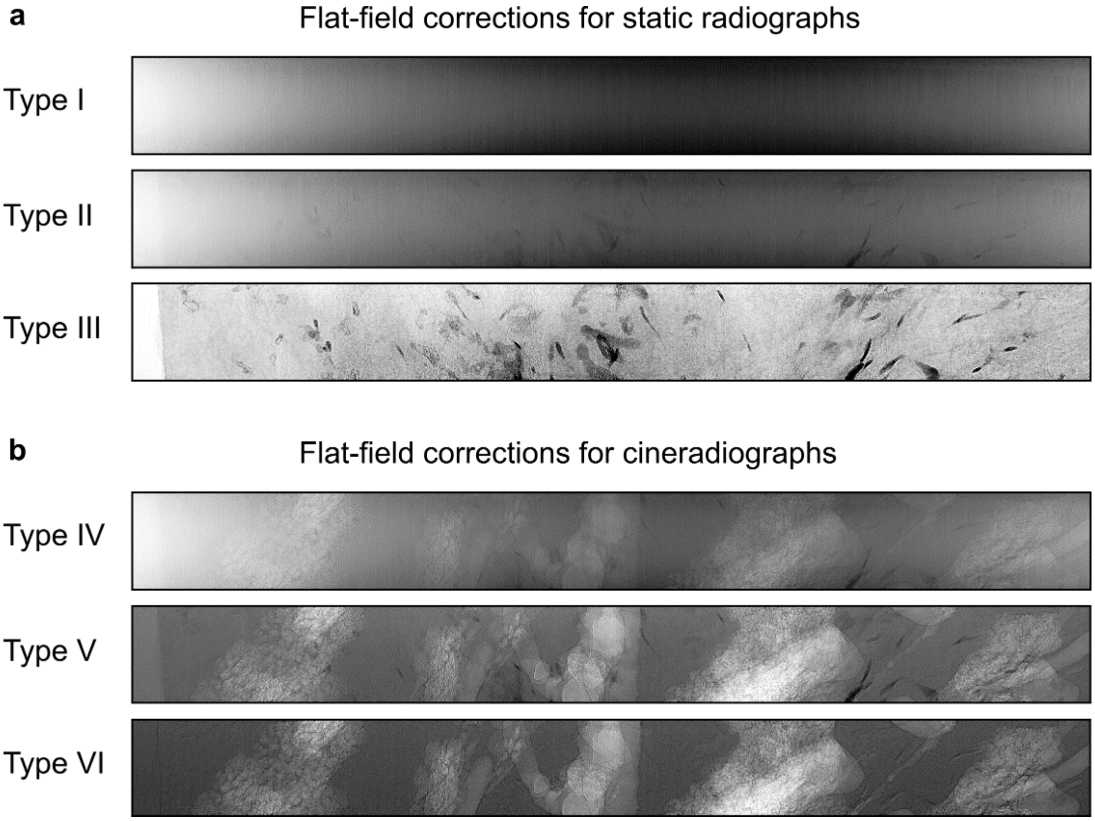

**Supplementary Fig. 4** Multiple flat-field corrections (FFCs) for enhancing in-tissue bubble contrast. The FFC outcomes are shown for **a**, static radiographs, including type I-III FFCs. **b**, For cineradiographs, the outcomes for type IV-V FFCs are shown. The definitions of all types of FFCs are given in Supplementary Table 1.

The static flat-field radiographs were measured before the cineradiography experiment. Type-I and type-II FFCs are classical X-ray absorption flatfield and are used to normalize the beam power profile as well as the detector’s inhomogeneities. As the flatfield references have been taken through the mounting

PP tube without the samples, this calculation removes the contribution of the air and of the mounting PP tube. These provide the absorption+propagation phase-contrast profile of agar and tissue + agar respectively, as well as of the sample jars (in PET). It has to be noticed that the residual vertical gradients in these pictures come from the vertical gradient in energy in the filtered white beam used on BM05 beamline. The average energy is slightly lower when going off the central beam, leading to higher absorption by the sample. Nevertheless, this effect is corrected by the other types of FFC described hereafter, as they are all integrating this gradient effect. Type-III FFC is used for imaging tissue microstructures while removing the local tomography effect, as has been demonstrated recently with hierarchical phase-contrast tomography(1) and to determine the beam position on the sample before continuous monitoring by cineradiography. Type-IV FFC is used to estimate the bubble volume by comparing with the absorption profile estimated using type-I and type-II FFCs. Type-V FFC is used to emphasize the bubbles by comparing the frames before and after bubble onset. The $I_{\mathrm{ref}}$ is calculated using the median intensity image of the cineradiographs collected before bubble onset normalized by the synchrotron current (the number of electrons in the storage ring). All the subsequent FFC or calculations are normalized by the synchrotron current at the exact time of each frame. The outcomes of these FFC schemes are shown in Supplementary Fig. 4.

#### S2.2. Bubble volume and growth quantification

The flat-field corrected phase-contrast cineradiographs contain signatures of the material components (air, agar, bubbles). To quantify the time-dependent bubble volume while accounting for the curved shape of

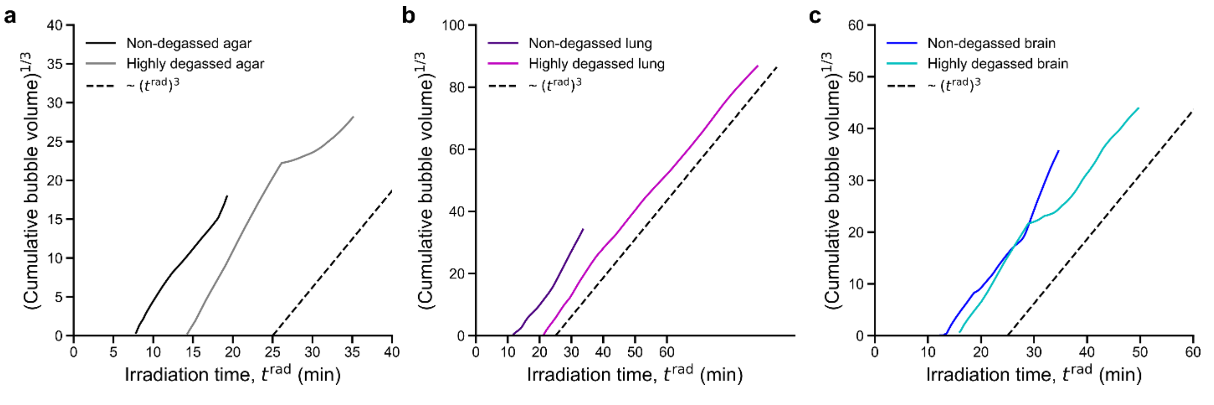

**Supplementary Fig. 5** The power-law behavior of bubble growth illustrated using the cubic root of the cumulative bubble volume over irradiation time for **a**, agar, **b**, lung slices (and agar), and **c**, brain slice (and agar) samples. The volumes are estimated within the field of view of the X-ray beam. Compared with the main text Fig. 3b, which show only a portion of the time trace after adjusting for bubble onset time in each case, **a**-**c** here shows the full duration of measurement (under X-ray irradiation). A reference curve is shown as a dashed black line to guide the eye for the trend-seeking.

the cylindrical sample container, we carried out a series of data processing to obtain the volume estimation of bubbles present in the field of view of the X-ray beam:

(i) Calculate the flat-field corrected radiographs ($I_{\mathrm{ff}}$) following type-I, II, IV, and V FFC.

(ii) Apply a median filter to remove phase fringes

(iii) Estimate the effective linear absorption coefficient ($\mu_{\mathrm{eff}}$).

(iv) Image segmentation of the projected areas ($\Omega_{\mathrm{bubble}}$) where the bubbles are present in $I_{\mathrm{ff}}$.

(v) Assuming a single material composition, we retrieve the bubble depth along the X-ray beam direction ($d$) is retrieved by inverting the Beer-Lambert law at each horizontal location $x$,

$$d(x) = -\frac{\log I_{\mathrm{ff}}}{\mu_{\mathrm{eff}}}. \tag{10}$$

(vi) Calculate the total bubble volume by integrating over all segmented spatial locations ($x, y \in \Omega_{\mathrm{bubble}}$).

$$V = \sum_{x,y} p_x p_y d(x), \tag{11}$$

where $p_x$ and $p_y$ are the pixel sizes of the radiograph (both are 25 µm in our case) along the two horizontal directions, respectively.

The bubble volume quantification procedure is performed over all cineradiographs to obtain the bubble size evolution over time with different samples and conditions. To avoid the granularity caused by the high temporal sampling of bubble growth, we used an integral measure, the cumulative bubble volume, to quantify the growth. To guide the eyes for visualization, we illustrate the growth using time versus the cubic root of the cumulative bubble volume, as shown in Supplementary Fig. 5, for reasons that will be clear in the data fitting result presented afterwards.

**Supplementary Table 2** Fitting results of bubble growth exponents within the first 10 mins after bubble onset.

| **Sample** | **Sample condition** | **Growth exponent for cumulative bubble volume** |
|---|---|---|
| Agar | non-degassed | 2.3 |
| Agar | highly degassed | 3.0 |
| Lungs slice + agar | non-degassed | 3.5 |

| | | |
|---|---|---|
| Lung slice + agar | highly degassed | 2.9 |
| Brain slice + agar | non-degassed | 2.7 |
| Brain slice + agar | highly degassed | 3.1 |

To quantify the bubble growth kinetics in the initial ~ 10 mins since the bubble onset for each case, we fit the cumulative bubble volume curves according to Eq. (2) in the main text using a least-squares approach to a power law model. To grasp the overall trend, we downsampled the data to timesteps of 0.5 mins before fitting. The fitted growth exponents are listed in Supplementary Table 2. The traces for highly degassed samples have more similar growth exponents around 3.0, while those for the non-degassed samples has a larger spread in the growth exponent, which depends on the sample content.

#### S3. X-ray dosimetry and temperature change estimation

X-ray dosimetry was carried out with a PTW Unidos E (T10021) dosimeter equipped with a TM31010 semiflex ionization chamber. The dosimeter was fixed at the sample position in air during measurement and an integrated incident dose rate of 36.83 Gy/s was obtained with the same X-ray beam condition as in radiography. The measurement outcome was used to estimate the temperature change within the sample during X-ray radiography. As the the synchrotron beam is decreasing over time and requires regular refills (every hour in the mode we used), the dose measurements have been considered for the average SR-current used for the whole experiment. Refills during the longest irradiations could explain some curve shape modifications for some samples.

**Supplementary Table 3** Estimated dose and temperature changes in samples at bubble onset time ($t^{bo}$).

| **Sample** | **Sample condition** | **Irradiation time (s)** | **Accumulated surface dose (kGy)** | **Temperature change (K) assuming 100% $H_2O$** | **Temperature change (K) assuming 100% EtOH** |
|---|---|---|---|---|---|
| Agar | non-degassed | 420 | 15.5 | 3.7 | 6.3 |
| Agar | highly degassed | 840 | 30.9 | 7.4 | 12.6 |
| Brain slice + agar | non-degassed | 780 | 28.7 | 6.9 | 11.7 |
| Brain slice + agar | highly degassed | 960 | 35.4 | 8.5 | 14.4 |

| | | | | | |
|---|---|---|---|---|---|
| Lung slice + agar | non-degassed | 600 | 22.1 | 5.3 | 9.0 |
| Lung slice + agar | highly degassed | 1260 | 46.4 | 11.1 | 18.9 |
| Mean value | non-degassed | 600 | 22.1 | 5.3 | 9 |
| Mean value | highly degassed | 1020 | 37.6 | 9 | 15.3 |

We assume no heat exchange between the tissue sample and the surrounding(13), which corresponds to an upper bound of the absorbed dose. Then, we relate the total incident energy ($Q$) to dose ($D$) and temperature change ($\Delta T$).

$$Q = Dm = c_p m \Delta T \,, \tag{12}$$

where $c_p$ is the specific heat of the system and $m$ is the weight of the sample, which we assume to be mostly solution. In the experiment, we measured the dose rate ($R_D$) of the X-ray beam. The accumulated surface dose may be calculated using the duration of X-ray irradiation ($t^{\mathrm{rad}}$). The estimated temperature change may then be obtained simply with,

$$\Delta T = \frac{R_D t^{\mathrm{rad}}}{c_p} . \tag{13}$$

Using Eq. (13), we estimated the temperature change from the start of the X-ray irradiation till bubble onset and tabulated the results in Supplementary Table 3. Due to its smaller heat capacity ($c_p^{\mathrm{H_2O}}$ = 4.184 J/(g·K), $c_p^{\mathrm{EtOH}}$ = 2.46 J/(g·K)), the temperature changes in solution assuming pure EtOH would be ~ 1.7 times higher than assuming pure $H_2O$.

### S4. Processing of micro-gas chromatography data

#### S4.1. Multi-module calibration

Before the synchrotron beamtime, we measured the chromatograms of distilled water, concentrated ethanol (96%, denatured), and agar mixed in 70% ethanol as background signals, as shown in Supplementary Fig. 6. These calibration measurements were conducted with glass bottles containing the corresponding solution placed in the same plastic housing (see Supplementary Fig. 2a) as the sample container was during phase-contrast cineradiography. During the calibration measurements, no X-ray irradiation was present to obtain a clean reference of the volatile species present in the mixture.

Within the detection limit, the chromatographic peaks representing air include $N_2$ and $O_2$, which are symmetric in shape. The peaks corresponding to $H_2O$ and EtOH both exhibit chromatographic tailing effects, likely due to their strong polarity. The concentrated ethanol was denatured when purchased and the denaturants occupy ~ 0.4% of ethanol, but no signatures of the denaturants have been found in the chromatograms among all used modules. The agar-ethanol mixture contains a more pronounced water peak due to its higher water concentration than pure ethanol. Since agar has a stable structure(14) and is non-volatile at room temperature, the chromatogram of the agar-ethanol mixture doesn't show any other peak from the corresponding ethanol chromatogram in all μGC modules.

##### S4.2. Null results for alternative chemical species

During the synchrotron experiments, we ran the four modules of the micro-gas chromatograph in parallel, where each module takes in a portion of gaseous eluent injected into the chromatograph(15). The

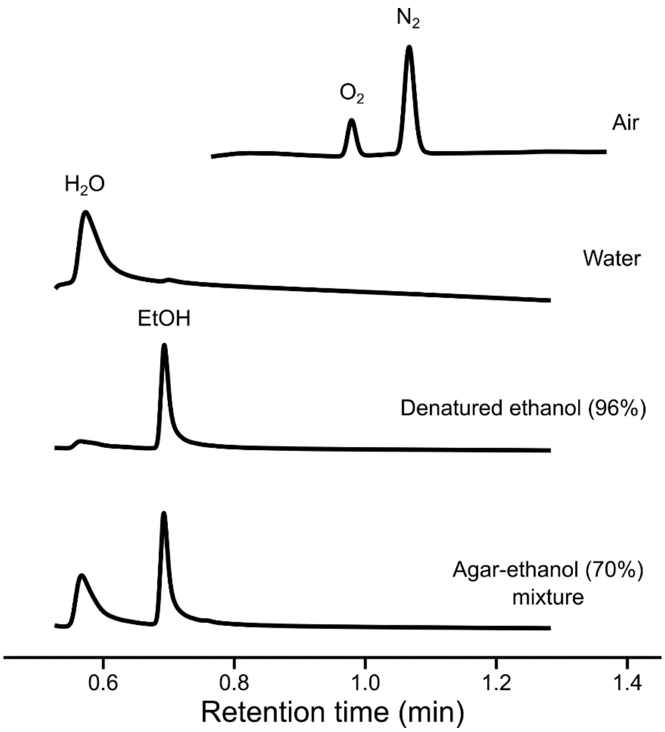

**Supplementary Fig. 6** Representative chromatograms for predominant volatile species measured by the micro-gas chromatograph APIX ChromPix2. The main components of air and dissolved gas ($N_2$ and $O_2$) are detected by the MS5A module, while the EtOH and $H_2O$ evaporated from the solutions are captured by the PDMS10 module.

concentration of the major components of air and dissolved gas ($N_2$ and $O_2$) were measured with the MS5A module, which detects lightweight gases (such as $N_2$, $O_2$, and $CH_4$). Concentrations of the solvent species ($H_2O$ and EtOH) were measured with the PDMS10 module (covering organic chemicals up to $C_6$). The other two modules (PDMS5 and PPU) were used to examine potential unexpected end products from photochemistry, such as those from the photodissociation and oxidation of agar and EtOH, just to name a few. The PPU module has a high sensitivity for $CO_2$ and $C_2H_2$, while the PDMS5 module detects heavier volatile organic chemicals from $C_6$ to $C_{10}$.

To time-correlate the operando gas chromatography signal with that obtained from phase-contrast cineradiography, we manually aligned the timestamp metadata of the corresponding data files. Since the cineradiography is acquired at a much faster rate (3 Hz) than gas chromatography (about 2.4 mins per round

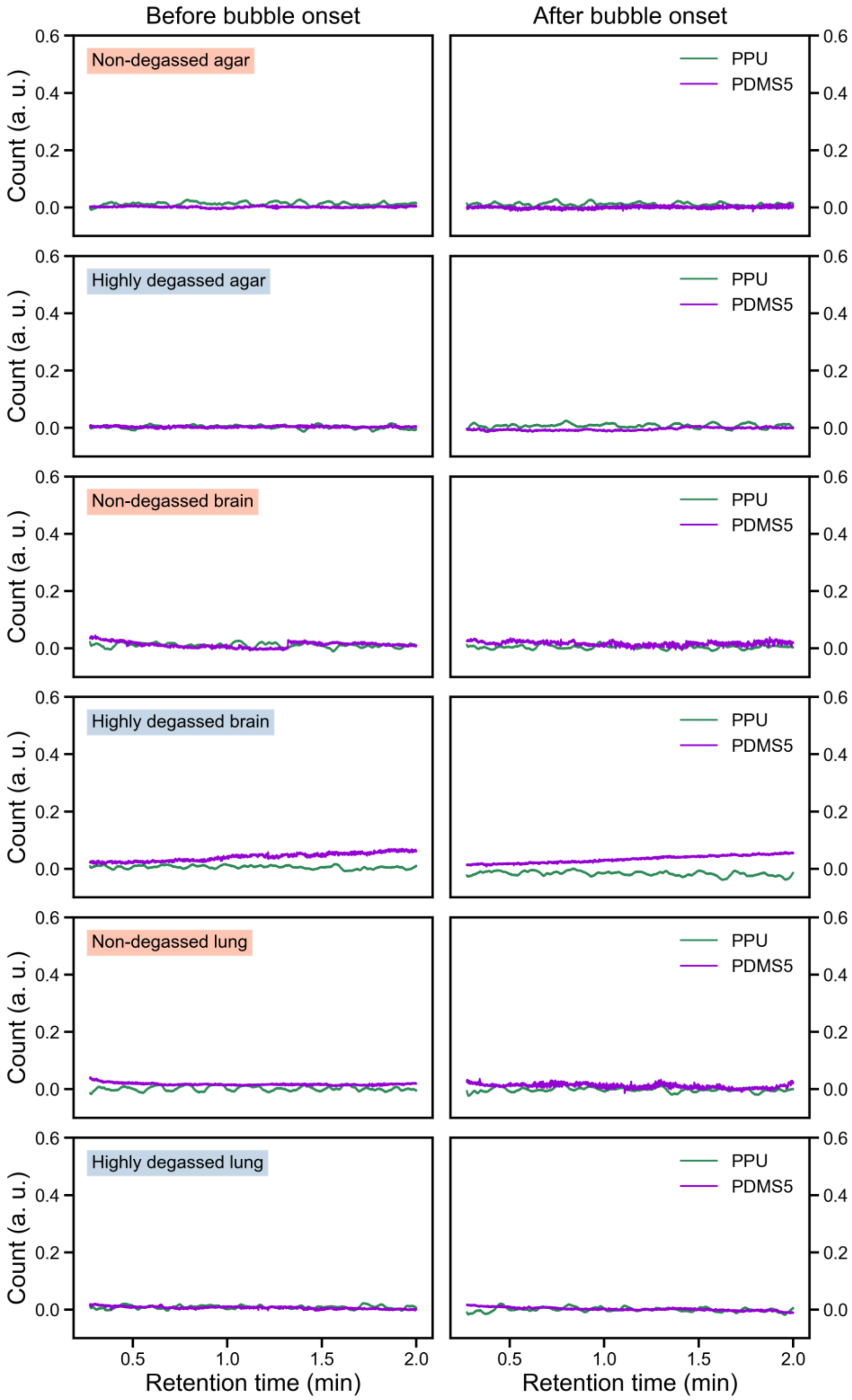

**Supplementary Fig. 7** Signals from PPU and PDMS5 modules of the micro-gas chromatograph. No salient feature has been observed for all tested samples in both modules, which aimed to detect alternative chemical products with higher molecular mass from radiation-matter interaction. In each case, the before- and after-spectrograms were measured at ~ 7 mins from bubble formation.

of elution), only approximate temporal alignment is possible. We processed the single-module readouts from the micro-gas chromatograph individually using area integration. For the two modules (MS5A and PDMS10) with identifiable chemical signals (see Supplementary Fig. 6), the relative concentrations were calculated using the ratio between the species. Since the experimental hutch, where the gas cylinder and chromatograph are placed, is not physically accessible during X-ray irradiation, and the experiments cannot be paused during timekeeping of the bubble dynamics, this data processing approach removes the slow drift from the carrier gas pressure regulators during experiments as well as the eventual gas flow differences between the samples as the closure of the system was not fully reproducible from one sample to the next one. Apart from the results shown in the Main Text Fig. 4, the results from the other two modules (PDMS5 and PPU) of the micro-gas chromatograph in other chemical sensitivity ranges do not contain anything noticeable, as shown in Supplementary Fig. 7.
